# Maximum Parsimony Interpretation of Chromatin Capture Experiments

**DOI:** 10.1101/533992

**Authors:** Dirar Homouz, Andrzej Kudlicki

## Abstract

We present a new approach to inferring global geometric state of chromatin from Hi-C data. Chromatin conformation capture techniques (3C, and its variants: 4C, 5C, Hi-C, etc.) probe the spatial structure of the genome by identifying physical contacts between genomic loci within the nuclear space. In whole-genome conformation capture (Hi-C) experiments, the signal can be interpreted as spatial proximity between genomic loci and physical distances can be estimated from the data. However, the results of these estimations suffer from internal geometric inconsistencies, notoriously violating the triangle inequality. Here we propose that the inconsistencies may be caused not by experimental artifacts but rather by a mixture of cells, each in one of several conformational states, contained in the sample. We have developed and implemented a graph-theoretic approach that identifies the properties of these subpopulations. We show that the geometrical conflicts in a standard yeast HiC dataset, can be explained by only a small number of homogeneous populations of cells (4 populations are sufficient to reconcile 95,000 most prominent impossible triangles, 8 populations can explain 375,000 top geometric conflicts). Finally, we analyze the functional annotations of genes differentially interacting between the populations, suggesting that each inferred subpopulation may be involved in a functionally different transcriptional program.

**Author Summary:** The global conformation of chromatin within a nucleus plays an important role in regulation of genes. The Hi-C technique can detect proximity between genomic loci, but attempts to use Hi-C data to infer the global conformation lead to hundreds of thousands of impossible geometries, violating the triangle inequality. To date, there was no explanation for this phenomenon. Here, we resolve these discrepancies by modeling the sample as a mixture of nuclei in several conformation states. We have developed a graph-theoretic approach to characterize the shape of chromatin in each of these states. In a real-life situation, as few as 4-5 discrete subpopulations can resolve all spatial inconsistencies in the data. Moreover, the results suggest that each subpopulation is associated with a functionally specific transcriptional program.

## Introduction

The three-dimensional organization of eukaryotic genome inside the nuclear space has been shown to play an important role in the regulation of transcription. In the last decade, our understanding of the genome organization has greatly progressed thanks to experimental techniques such as chromosome conformation capture (3C), 4C, 5C, 6C, ChIA-PET, and HiC [1–6]. In these methods, three-dimensional contacts between different parts of the DNA are captured by ligation, and characterized, typically by sequencing and mapping to their genomic loci. The contacts provide information on the spatial organization of the genome. Recent developments in these technologies have allowed high-resolution mapping of the interactions within in the entire genome.

The spatial, Euclidean distances between interacting loci can be estimated from chromatin capture data [7, 8], and may, in turn, be used for constructing three-dimensional models of the genome [8].

We have observed that 3-D models constructed from DNA interaction data interpreted as distances within a single conformation, may lead to impossible geometries. Specifically, in a haploid cell, a conformation is not possible in which *locus C* is close to *locus A* and to *locus B*, but the Euclidean distance between the loci *A* and *B* is large:

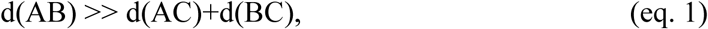

which leads to an impossible triangle (d() denotes the physical, Euclidean distance between two loci in the nuclear space). Such a situation corresponds to strong 3C signal for the AC and BC interactions, but few or no reads that would correspond to interaction between A and B. When using the standard formula to estimate the Euclidean distance between loci from Hi-C reads in the yeast HiC data of [6], we find large numbers of such impossible triangles. Observation of such apparently impossible geometries may be attributed to a range of possible causes, including inaccuracy of the formula, noise in the data, systematic errors such as sequencing bias, and others [9], including experimental errors [10]. Another possible explanation is that the chromatin conformation is not rigid but constantly changing and that the HiC experiments represent not one conformation but rather an entire ensemble of states accessible by small, thermal-like motions.

Such data-driven models implicitly assume that the population properties of cells used in the experiment are uniform with respect to the chromatin conformation; however, this is not necessarily true, as it is known from fluorescent microscopy that the chromatin structure may vary between cells [11]. The inability to appreciate the dynamics and variability of chromatin states has been indicated as the main drawback of Hi-C based methods [12], triggering research into developing single-cell Hi-C experimental techniques.

Here, we take this proposition to the opposite extreme: rather than considering a continuous ensemble of possible chromatin conformations, we investigate whether it is possible that the measured HiC signal is produced by a small number of discrete, rigid conformations of DNA and how many such conformations would be required, and what are their properties. To this end, we developed a graph-theoretic approach to this problem, that is presented below, along with and the resulting characterization of such postulated rigid states.

## Results

### Characterization of the geometric conflicts

As our primary dataset, we use the standard yeast data of Duan et al [6], providing DNA contact information with kilobase resolution for a model haploid genome. The experiment probed chromatin interactions for all pairs of HINDIII restriction enzyme target loci in the yeast genome. To convert read counts to approximate physical distances, we use the formula derived by [7]:

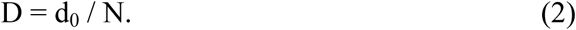

The value of d_0_ has been estimated by [6] as 155,000 nanometers, however, in the present considerations the numeric value is never used in the calculations, it will cancel out as long as it remains approximately constant.

To characterize the conflicts, we introduce a working definition of the triangle inequality [Eq. 3]. We call a triangle ABC (AB being the longest side) “*impossible”* when the estimated distances satisfy the following condition:

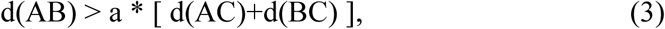

for a given a > 1.

We applied equation (3) to all triples of HINDIII loci in the HINDIII dataset of ([6]). As a result, for values of *a* from 1.3 to 2, we obtain between 30953 and 664101 conflicts (impossible triangles) in the genome that affect the majority of HINDIII sites in the genome (See column 2 and 3 of Table 1).

**Table 1:** Characterization of geometric conflicts (violations of triangle inequality) for the yeast dataset of Duan et al, at different thresholds of conflict definition *a*.

| $a$ | Conflicts | HINDIII loci involved in conflicts | In coding sequence | within 500bp from TSS | HINDIII loci acting as mid loci | Mid loci in coding sequences | Mid loci within 500bp from TSS |
| --- | --- | --- | --- | --- | --- | --- | --- |
| 1.3 | 664101 | 3476 | 1338 | 460 | 2810 | 1074 | 360 |
| 1.4 | 374735 | 3442 | 1323 | 455 | 2712 | 1032 | 348 |
| 1.7 | 95370 | 3375 | 1298 | 446 | 2326 | 889 | 295 |
| 2.0 | 30953 | 3247 | 1246 | 428 | 1834 | 700 | 228 |

Since the 3D structure of the genome is associated with transcriptional regulation, we next tested whether there is a dependence between HINDIII site involvement in conflicts and its association with coding sequence or transcription start sites. Since a vast majority of HINDIII loci in the yeast genome are involved in at least one of the geometrical conflicts, the global enrichments are not highly significant. An impossible triangle (as defined by Eq 1 and Figure 1) consists however of three HiC loci, the “promiscuous”, mid-site C that is in contact with both other two sites, and the “exclusive” A and B ends that show Hi-C signal only with C, but not with each other. While most HINDIII loci show some involvement in the conflicts, only a fraction of them assume the role of the promiscuous, “C” vertex of the triangle. Still, the fraction of “mid” C-loci to all loci in contacts does not change significantly when only loci near TSS, or loci in coding sequence are considered.

**Fig 1:**
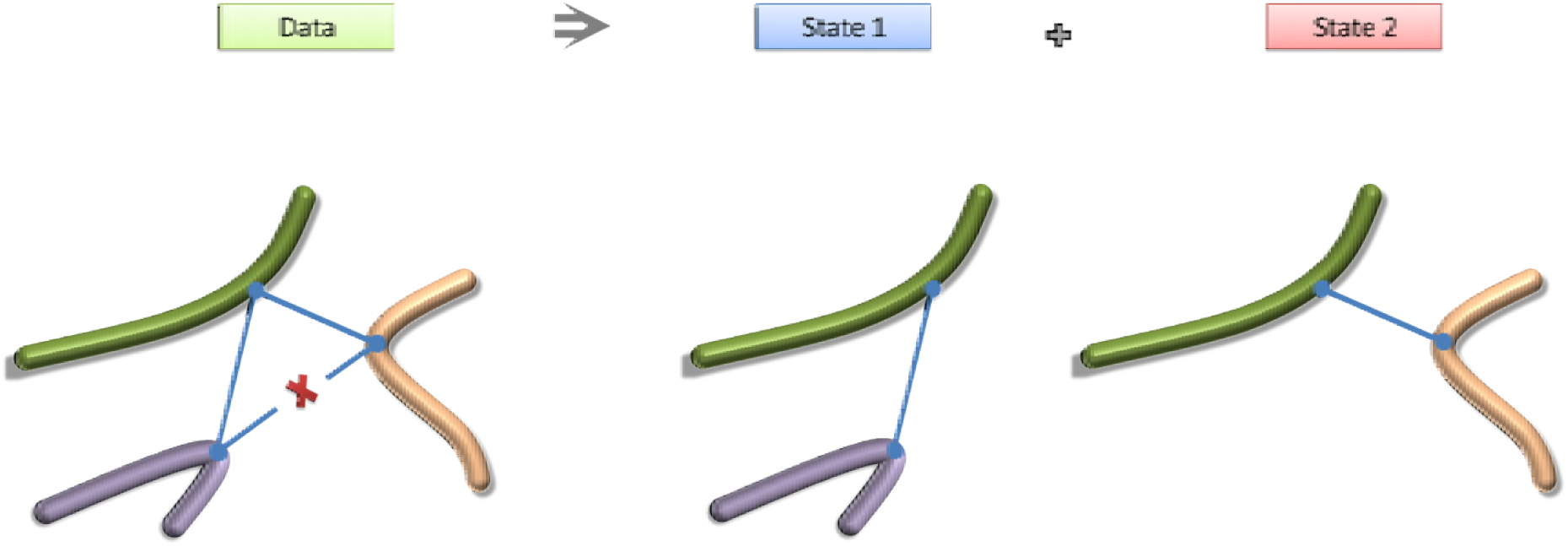
A schematic representation of three loci in a haploid genome, forming an impossible triangle. If tight DNA contacts (blue lines) are observed for two pairs of loci, but no signal is present for the third one (x), then the two observed interactions cannot coexist in one type of haploid cell and must represent distinct chromatin states.

## Resolving the spatial conflicts

### The mixed-state hypothesis

The fact that a large number of geometric inconsistencies are present in Hi-C data makes it a crucial problem in interpreting chromatin capture 3C experiments and calls for identifying a solution to the issue. The existence of an impossible triangle may be explained by a range of factors, such as experimental error, or limited applicability of Eq. (2). It is also possible that stochastic motions occasionally bring loci A and C or loci B and C together, but for some reasons loci A and B are never in close spatial proximity.

Here, we introduce an alternative hypothesis and verify that it is consistent with the data. We propose that the Hi-C data may result from a small number of discrete conformations of genomic DNA, each conformation corresponding to a specific subpopulation of cells within the experimental sample. An “impossible triangle” inferred from the data corresponds to two interactions (AC and BC) whose coexistence is ruled out due to lacking evidence of proximity between A and B. Our working hypothesis is rooted in the proposition that AC and BC indeed do <u>not</u> coexist in the same cells; as some cells have only the AC interaction while others only BC. The concept of disentangling the population-averaged measurements is presented graphically in Figure 1.

While resolving one geometrical conflict requires two distinct subpopulations of cells in the experimental sample, one might expect that the number of subpopulations needed to explain the thousands of conflicts in the data would be very large, suggesting that the approach is not practical at all. However, just one pair of subpopulations may be enough for explaining more conflicts. For example, if AC is incompatible with BC, and XZ cannot coexist with YZ, it is possible within our framework that AC and XZ exist in subpopulation I, while BC and YZ coexist in subpopulation II. Below we introduce and implement a graph-theoretic approach to determine and characterize the minimal number of subpopulations required to reconcile all the conflicts found in the data.

### Globally resolving conflicts in Hi-C contacts

The list of all observed conflicts is used to create a graph G = (V, E) where each vertex in V represents a contact (a pair of interacting sites; note that the graph is dual to a graph representing genomic loci as vertices and interactions as edges). Two vertices are connected by an edge in E if they are in conflict, that is the two interactions cannot coexist in the same homogeneous subpopulation of cells (Figure 2). The problem of finding the minimal number of subpopulations that can reconcile the experimental data is equivalent to coloring the vertices of the graph G such that two vertices with the same color must not be connected with an edge (two interactions occurring in the same subpopulation must not be incompatible due to forming an impossible triangle). This means that two conflicted contacts will be colored differently and thus always belong to different colors (or states of the genome). A graph-coloring algorithm will find the chromatic number of the graph, i.e. minimum number of colors for a graph and will color the vertices according to this minimum. Coloring of vertices of a graph is a classical problem in mathematics and computer science. The graph coloring algorithm we used is based on the column generation principle. The graph coloring algorithm that we used here is a modification of the heuristic approach of M. Trick [13, 14], see Methods; the algorithm is better suited to large graphs than exact methods such as of DSATUR [15].

**Figure 2:**
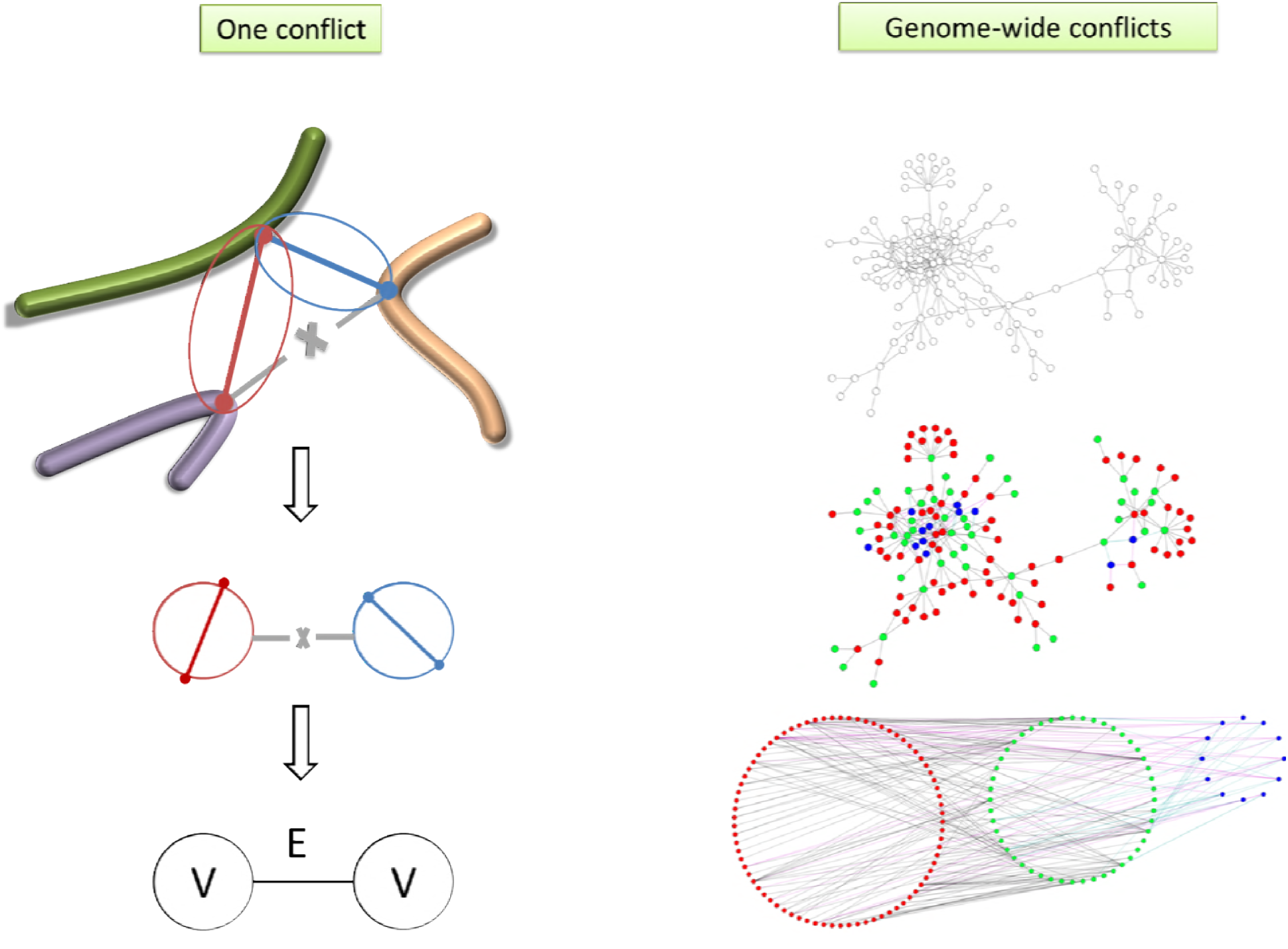
Resolving contact conflicts. Left: Generating the conflict graph, where the vertices are contacts and edges are conflicts. Right: Assuming a mixed population of cells in the sample, we resolve the conflict by coloring the (V,E) conflict graph.

Figure 2 summarizes the approach used to globally resolve the contact conflicts: defining the graph of conflicts, the coloring approach and the interpretation of the colors. We applied the coloring algorithm to the conflict graphs generated from the yeast Hi-C dataset, with several values of the cutoff parameter a between 1.3 and 2.0. The results agree with our expectation that the large numbers of conflicts can be reconciled with only several homogeneous conformations contained in the sample.

The results are summarized in Table 2, and an example coloring of a subset of the interaction graph for a=2.0 is shown in Figure 2 (right panel).

**TABLE 2:** Minimum numbers of colors (genome states) needed to reconcile the yeast HiC data, depending on the threshold of *a* in conflict definition (Eq.1).

| a | Pairs in conflicts | N colors | Locus-pairs with conflicting interactions of at most this many colors |  |  |  |  |  |  |  |  |  |
| --- | --- | --- | --- | --- | --- | --- | --- | --- | --- | --- | --- | --- |
|  |  |  | 1 | 2 | 3 | 4 | 5 | 6 | 7 | 8 | 9 | 10 |
| 1.3 | 86781 | 10 | 21702 | 40809 | 58248 | 76832 | 83164 | 85957 | 86462 | 86703 | 86763 | 86781 |
| 1.4 | 62093 | 8 | 17610 | 32010 | 48377 | 57581 | 61307 | 61971 | 62070 | 62093 |  |  |
| 1.7 | 30085 | 5 | 10255 | 21636 | 28862 | 30050 | 30085 |  |  |  |  |  |
| 2.0 | 16551 | 4 | 7057 | 14734 | 16528 | 16551 |  |  |  |  |  |  |
| Rand | 22006 | 22 | 7420 | 10863 | 13248 | 15096 | 16652 | 17959 | 19053 | 19909 | 20547 | 21004 |

The result confirms that the HiC data of ([6]) can indeed be reconciled as a product of a small number of subpopulations in the experimental sample. Specifically, using the threshold *a*=2.0, we can demonstrate that as few as 4 subpopulations of cells are sufficient to explain all the 30953 conflicts in the data. Moreover, even for low *a* there are only very few loci whose interactions require more than 5 colors. This observation may suggest that the majority of the conflicts are caused by the presence of only 3-4 rigid, homogeneous subpopulations and the remaining conflicts are only a very small fraction of the total and are caused by some kind of experimental artifact, possibly sequencing bias. It is important to note, that by modeling the sample as a mixture of only a few populations, we reduce the number of geometric conflicts from hundreds of thousands to zero, which constitutes an obvious improvement over previous approaches to infer the global geometry of the genome.

Finally, we demonstrate that the distribution properties of the network of interactions between the states (“colors”) is not consistent with a random distribution of conflicts. To this end, we randomized the assignment of edges in the conflicts graph (to represent random conflicts between existing 3C interactions). The result is drastically different (see bottom row of Table 2), suggesting that the conflicts do not arise from measurement errors, but from actual states of the shape of the genome.

In most cases, there are multiple solutions for coloring a graph using the minimal number of colors (See Fig 3a). To test the stability, or similarity between alternative solutions, we ran the coloring algorithm 20 times, randomly re-ordering the list of graph nodes on its input. In each case, the algorithm produced the same number of colors, but the assignments of colors to interactions was different in each solution. To test the consistency of coloring (or assess the similarity between the results from different random seeds), we assessed how many pairs of interactions that were in the same color in one solution remained of the same color in other solutions. The results are summarized in Figure 3b and in Supplementary Table ST1, and demonstrate that although not identical, the solutions have a significant degree of similarity, distinguishing them from random assignment of states. In other words, there is a significant global coordination between contacts across the entire genome; the identity of interactions attributable to each state is genuine.

**FIGURE 3.**
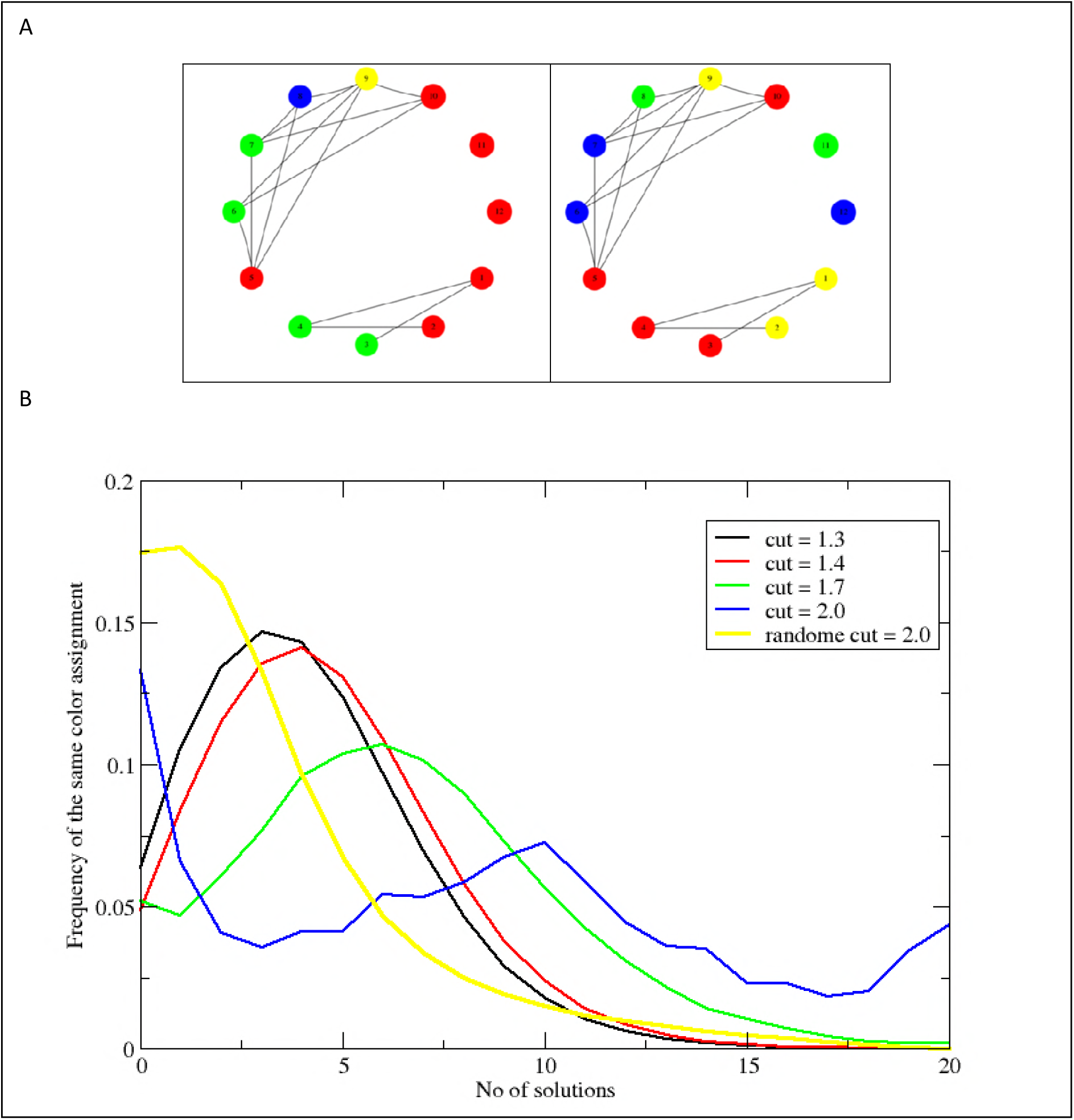
Alternative coloring solutions. A: Illustration of different colorings of the same conflict graph. B: Consistency between alternative coloring solutions in yeast HiC data. The figure shows the histograms of the number of similar color assignments among 20 possible coloring solutions. The histogram for the HiC contact conflict graph shows reasonable stability as evident by the fact that the histogram has its peak away from zero (a=1.3,1.4, and 1.7). On the other hand, a random graph has its peak at zero which means that the coloring solution is less stable than real data. Finally, we see that the graph loses its coupling at high a, causing the peak at zero for a=2.

### Functional characterization of the sub-populations

We have demonstrated that the apparent conflicts observer in Hi-C data can be explained by a small number of homogeneous subpopulations of cells contained in the sample. Statistical considerations presented above suggest that at least to some extent the subpopulations are genuine and reflect biologically relevant coordination between contacts in different parts of the genome. It has been demonstrated that a correspondence exists between spatial organization of the yeast genome and transcriptional regulation of genes: genes in interacting genomic loci are often coexpressed and share the same functional annotations [16]; the dependence also exists in other species [17–19]. If the chromatin states of cell subpopulations are indeed real, they should be subject to evolutionary pressure, and thus functionally relevant. In order to understand the biological significance of these states, we analyzed the distribution of groups of genes with different functions among these states. To confirm the functional role of the different conformational states we calculated the enrichment of different GO-slim term within the 6 states and compared with that of randomly created states. As it can be seen from Figure (4), many groups of genes tend to behave differently in different conformational states confirming the dynamical role that the genome conformation plays in regulating the functions of genes. The figure also shows that the number of significantly depleted or enriched groups of genes in the 6 conformational states is much higher than that of 6 random states confirming the nonrandom nature of these 6 genome confirmations.

The same dynamic behavior of these conformational states is also observed in different cell cycle phases [20] and metabolic cycle clusters [21]. The data for the cell cycles is shown in the inset of Fig. 4. The results suggest that, especially during the cell cycle, specific inferred conformational states of the genome bring together genes that are active in specific phases of the cycle.

**Figure 4:**
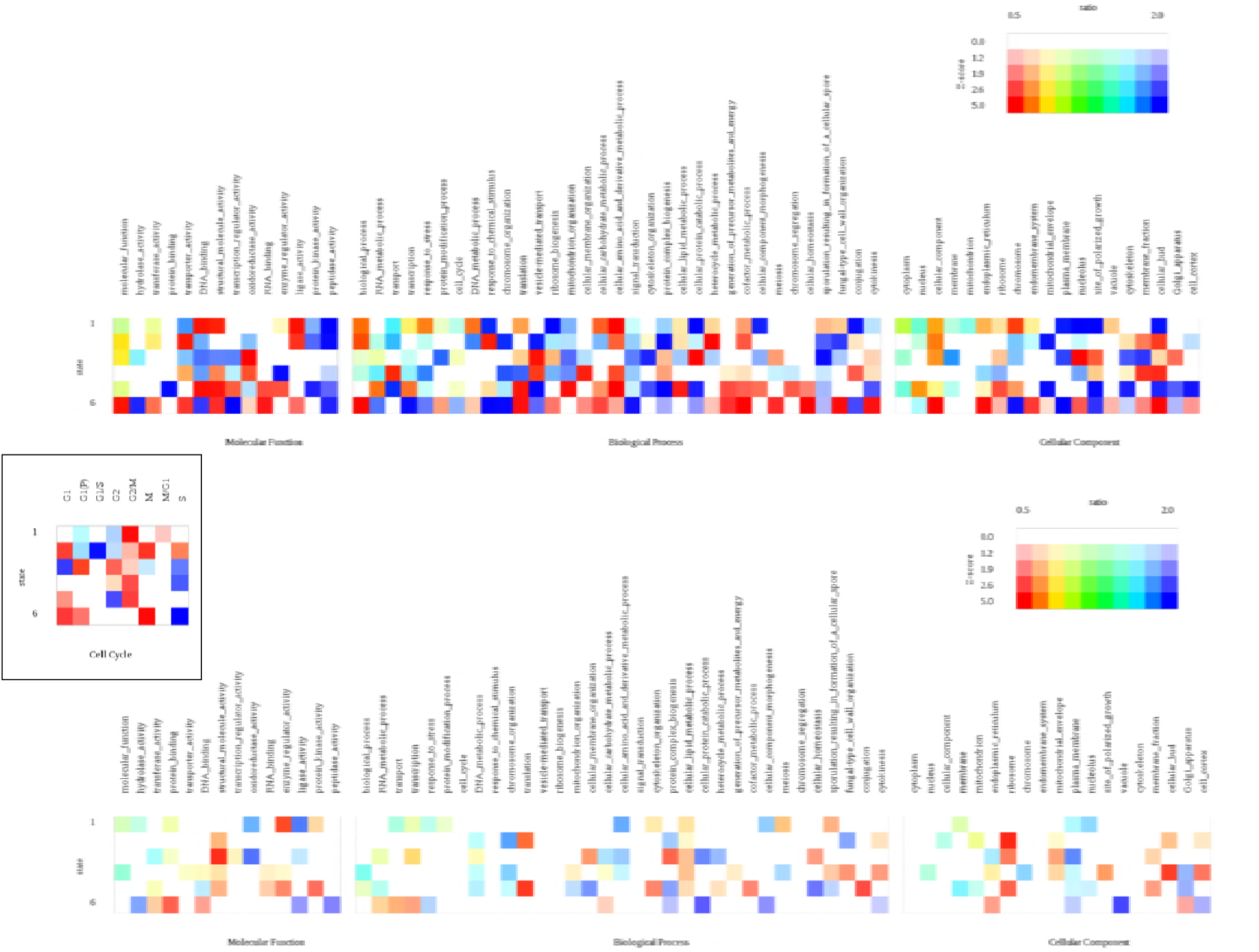
Analysis of functional annotation (significant GO terms) of genes whose loci interact in each of 5 inferred conformational states of the genome (top), compared to similar analysis for locus pairs with randomly assigned states (bottom). Inset: enrichments of genes associated with specific phases of the cell cycle.

## Conclusions

In this work, we have presented the first technique to globally reconcile geometric inconsistencies observed in HiC data. Our approach may lead to new way of understanding the dynamics of genome conformations and its interplay with transcriptional programs. We show that such apparent geometric conflicts may be explained by a mixture of states present in the experimental sample. We have designed a graph-theoretic approach to determine the number of states needed to reconcile the data. Our implementation of the approach allows to find example solutions for genomic loci and their interaction that is limited to a specific conformational state of the genome. By applying the method to a yeast HiC dataset, we have shown that HiC data can be interpreted as a mixture of a small number of homogeneous states (we do not conclude that it *is* such a mixture, but that such interpretation is consistent with the data). Coordination between pairs of loci interacting in these states suggests the states may be biologically relevant/significant. The biological significance is supported by analysis of the functional categories of genes interacting in each thus defined state. The functional annotations are non-random, which is consistent with the hypothesis that the mixture of states actually exists, and that different states may be associated with the cells executing specific transcriptional programs. The possibility of global, coordinated changes in chromatin conformation may have consequences not only for transcriptional programming, but also for genome stability genome instability and DNA repair [22]; such interplay may be confirmed by correlating the inferred states with DNA damage patterns studied at high resolution [23, 24]. Finally, the approach may be generalized to use also with diploid nuclei and applied to studying conformation dynamics of the human genome. It is also important to note that our graph-coloring approach is not limited to distances and conflicts defined by Equations (2) and (3) but it is a general framework that can be used with any method of assessing the distance and its error from the number of reads in the HiC dataset. Our simple, computational approach provides an alternative to technically challenging single-cell chromatin capture experiments.

## Methods

### Identifying conflicts in 4C contacts

The Hi-C (referred to as “4C” by [8]) yeast dataset consists of a list of genomic loci of different captured DNA fragments. The genomic positions for the two ends of each fragment are provided in addition to the count frequency of each fragment. The count frequency for a captured fragment represents the contact probability between the two sites connected by the fragment. The frequency is also related to the distance between contacted sites. The contact frequency can be converted into a distance assuming that this frequency is equivalent to that of a polymer packing problem, yielding an approximately inversely proportional dependence. Thus, for each observed contact, we can estimate the Euclidean distance between the two ends of that contact.

In order to investigate whether the 4C contact data for yeast represents a static conformation or multiple conformational states, we look for conflicts between triplets of interacted sites. In the data under consideration, there are many triplets of contacted sites that would form triangles. Assuming one static conformation of the yeast genome, the observed distances in each triangle should obey the triangle inequality. On the other hand, the presence of different conformational states in the data will manifest itself in triplets that violate the triangle inequality (Figure 1). Those triplets that violate triangle inequality represent the contact conflicts that we seek to analyze. The conflicts are characterized by impossible triangles where one of the edges is larger than the total length of the other two or that one of the edges of the triangle is missing. The missing contacts are assigned a frequency of 4 (The minimum reported frequency is 5).

### Resolving conflicts in 4C contacts

The list of all observed conflicts is used to create a dual graph G = (V, E) where each vertex in V represents a contact (two interacting sites). Two vertices are connected by an edge in E if they are in conflict (Figure 2). In order to resolve these conflicts, we utilize the graph coloring techniques from the graph theory. The graph coloring scheme used here is “vertex coloring” where two connected vertices are colored (labeled) differently. This means that two conflicted contacts will be colored differently and thus belong to different colors (or states). A graph coloring algorithm will find the minimum number of colors for a graph and color the vertices according to this minimum. The graph coloring algorithm used here is a modified heuristic argument of [13, 14]. We used Michael Trick’s modified heuristic approach [13, 14] rather than other popular, exact methods such as of DSATUR [15], since it is significantly faster while producing equivalent results. Running the coloring code of [14] with the large graphs is memory-intensive. For this reason, we optimized the code by modifying the memory management and replacing 4-byte integers with 1-byte where adequate; the resulting implementation was able to process the graph for the cutoff factor a=1.3 using less than 26GB of RAM. The algorithm has been validated to produce correct coloring results on randomly colored graphs (Supplementary Table ST2). Figure 2 summarizes the scheme used to resolve the contact conflicts.

## Acknowledgment

We thank Gang Chen for assistance with code optimization and testing. The research was partly supported by Clinical and Translational Science Award from the National Center for Advancing Translational Sciences UL1TR000071 and NIH grant GM112131.

## Supporting Information

**Supplementary Table ST1**: Stability of color assignment in HiC conflict data. See discussion in text and Figure 3b.

**Supplementary Table ST2**. Validation of the coloring algorithm. The program correctly reconstructs coloring of a random graph 96%-100% of the time in situations similar to the HiC conflict graphs.

## Bibliography

1. Lieberman-Aiden E, van Berkum NL, Williams L, Imakaev M, Ragoczy T, Telling A, et al. Comprehensive Mapping of Long-Range Interactions Reveals Folding Principles of the Human Genome. Science. 2009;326(5950):289–93. PubMed PMID: ISI:000270599500043.

2. Dekker J, Rippe K, Dekker M, Kleckner N. Capturing chromosome conformation. Science. 2002;295(5558):1306–11. PubMed PMID: ISI:000173926000047.

3. Simonis M, Klous P, Splinter E, Moshkin Y, Willemsen R, de Wit E, et al. Nuclear organization of active and inactive chromatin domains uncovered by chromosome conformation capture-on-chip (4C). Nature Genetics. 2006;38(11):1348–54. PubMed PMID: ISI:000241592700026.

4. Zhao Z, Tavoosidana G, Sjolinder M, Gondor A, Mariano P, Wang S, et al. Circular chromosome conformation capture (4C) uncovers extensive networks of epigenetically regulated intra- and interchromosomal interactions. Nature Genetics. 2006;38(11):1341–7. PubMed PMID: ISI:000241592700025.

5. Dekker J. The three 'C's of chromosome conformation capture: controls, controls, controls. Nat Methods. 2006;3(1):17–21. doi: 10.1038/nmeth823. PubMed PMID: ISI:000234528000011.

6. Duan Z, Andronescu M, Schutz K, McIlwain S, Kim YJ, Lee C, et al. A three-dimensional model of the yeast genome. Nature. 465(7296):363–7. doi: 10.1038/nature08973. PubMed PMID: ISI:000277829200044.

7. Bystricky K, Heun P, Gehlen L, Langowski J, Gasser SM. Long-range compaction and flexibility of interphase chromatin in budding yeast analyzed by high-resolution imaging techniques. Proceedings of the National Academy of Sciences of the United States of America. 2004;101(47):16495–500. PubMed PMID: ISI:000225347400023.

8. Duan Z, Andronescu M, Schutz K, McIlwain S, Kim YJ, Lee C, et al. A three-dimensional model of the yeast genome. Nature. 2010;465(7296):363–7. doi: Doi 10.1038/Nature08973. PubMed PMID: ISI:000277829200044.

9. Duggal G, Patro R, Sefer E, Wang H, Filippova D, Khuller S, et al. Resolving spatial inconsistencies in chromosome conformation measurements. Algorithms Mol Biol. 2013;8(1):8. doi: 10.1186/1748-7188-8-8. PubMed PMID: 23497444; PubMed Central PMCID: PMCPMC3655033.

10. Duggal G, Patro R, Sefer E, Wang H, Filippova D, Khuller S, et al. Resolving spatial inconsistencies in chromosome conformation measurements. Algorithms for Molecular Biology. 2013;8. PubMed PMID: ISI:000319000700001.

11. Bolzer A, Kreth G, Solovei I, Koehler D, Saracoglu K, Fauth C, et al. Three-dimensional maps of all chromosomes in human male fibroblast nuclei and prometaphase rosettes. Plos Biology. 2005;3(5):826–42. PubMed PMID: ISI:000229125400012.

12. de Wit E, de Laat W. A decade of 3C technologies: insights into nuclear organization. Genes Dev. 26(1):11–24. Epub 2012/01/05. doi: 26/1/11 [pii]10.1101/gad.179804.111. PubMed PMID: 22215806; PubMed Central PMCID: PMC3258961.

13. Mehrotra A, Trick MA. A column generation approach for graph coloring. informs Journal on Computing. 1996;8(4):344–54.

14. Trick MA. Available from: http://mat.gsia.cmu.edu/COLOR/color.html.

15. Brélaz D. New methods to color the vertices of a graph. Communications of the ACM. 1979;22(4):251–6.

16. Homouz D, Kudlicki A. The 3D Organization of the Yeast Genome Correlates with Co-Expression and Reflects Functional Relations between Genes. Plos One. 2013;8(1):e54699. doi: 10.1371/journal.pone.0054699.

17. Liu C, Wang C, Wang G, Becker C, Zaidem M, Weigel D. Genome-wide analysis of chromatin packing in Arabidopsis thaliana at single-gene resolution. Genome Res. 2016;26(8):1057–68. doi: 10.1101/gr.204032.116. PubMed PMID: 27225844; PubMed Central PMCID: PMCPMC4971768.

18. Dong X, Li C, Chen Y, Ding G, Li Y. Human transcriptional interactome of chromatin contribute to gene co-expression. BMC Genomics. 2010;11:704. doi: 10.1186/1471-2164-11-704. PubMed PMID: 21156067; PubMed Central PMCID: PMCPMC3053592.

19. Babaei S, Mahfouz A, Hulsman M, Lelieveldt BP, de Ridder J, Reinders M. Hi-C Chromatin Interaction Networks Predict Co-expression in the Mouse Cortex. PLoS Comput Biol. 2015;11(5):e1004221. doi: 10.1371/journal.pcbi.1004221. PubMed PMID: 25965262; PubMed Central PMCID: PMCPMC4429121.

20. Rowicka M, Kudlicki A, Tu BP, Otwinowski Z. High-resolution timing of cell cycle-regulated gene expression. Proceedings of the National Academy of Sciences of the United States of America. 2007;104(43):16892–7. doi: Doi 10.1073/pnas.0706022104. PubMed PMID: ISI:000250487600032.

21. Tu BP, Kudlicki A, Rowicka M, McKnight SL. Logic of the yeast metabolic cycle: Temporal compartmentalization of cellular processes. Science. 2005;310(5751):1152–8. doi: 10.1126/science.1120499. PubMed PMID: ISI:000233437300037.

22. Aymard F, Aguirrebengoa M, Guillou E, Javierre BM, Bugler B, Arnould C, et al. Genome-wide mapping of long-range contacts unveils clustering of DNA double-strand breaks at damaged active genes. Nature structural & molecular biology. 2017;24(4):353–61. Epub 2017/03/07. doi: 10.1038/nsmb.3387. PubMed PMID: 28263325; PubMed Central PMCID: PMCPMC5385132.

23. Biernacka A, Zhu Y, Skrzypczak M, Forey R, Pardo B, Grzelak M, et al. i-BLESS is an ultra-sensitive method for detection of DNA double-strand breaks. Commun Biol. 2018;1:181. Epub 2018/11/06. doi: 10.1038/s42003-018-0165-9. PubMed PMID: 30393778; PubMed Central PMCID: PMCPMC6208412.

24. Crosetto N, Mitra A, Silva MJ, Bienko M, Dojer N, Wang Q, et al. Nucleotide-resolution DNA double-strand break mapping by next-generation sequencing. Nat Methods. 2013;10(4):361–5. Epub 2013/03/19. doi: 10.1038/nmeth.2408. PubMed PMID: 23503052; PubMed Central PMCID: PMCPMC3651036.

